# Brief Report: The Virucidal Efficacy of Oral Rinse Components Against SARS-CoV-2 In Vitro

**DOI:** 10.1101/2020.11.13.381079

**Authors:** Evelina Statkute, Anzelika Rubina, Valerie B O’Donnell, David W. Thomas, Richard J. Stanton

## Abstract

The ability of widely-available mouthwashes to inactivate SARS-CoV-2 in vitro was tested using a protocol capable of detecting a 5-log10 reduction in infectivity, under conditions mimicking the naso/oropharynx. During a 30 second exposure, two rinses containing cetylpyridinium-chloride and a third with ethanol/ethyl lauroyl arginate eliminated live virus to EN14476 standards (>4-log10 reduction), while others with ethanol/essential oils and povidone-iodine (PVP-I) eliminated virus by 2-3-log10. Chlorhexidine or ethanol alone displayed little or no ability to inactivate virus. Studies are warranted to determine whether these formulations can inactivate virus in the human oropharynx *in vivo*, and whether this might impact transmission risk.

## Background

The lipid membranes of enveloped viruses are sensitive to disruption by lipidomimetic agents and surfactants. Thus, we hypothesised that the SARS-CoV-2 virus would be susceptible to inactivation by components in widely available mouthwashes, such as ethanol/essential oils, cetylpyridinium chloride (CPC) and povidone-iodine (PVP-I) [1]. Indeed, mouthwashes have been empirically employed in outbreaks in China during the current pandemic [2]. Two recent studies demonstrated that this approach can work *in vitro* under conditions that mimic nasal/oral passages. First, several formulations including dequalinium/benzalkonium chloride, PVP-I and ethanol/essential oils reduced SARS-CoV-2 infectivity *in vitro* by up to 3-log10 [3]. Second, infectivity of the closely related HCoV-229E coronavirus was reduced by 3-4-log10 using several agents including CPC, ethanol/essential oils and PVP-I [4]. Inactivation of HCoV-229E by >3-log10 by CPC at 0.07% was also shown in a recent preprint [5]. So far, only one of the products tested (Listerine Antiseptic, combining 26.9% alcohol with essential oils) achieved the 4-log10 kill required to pass EN14476 as a virucidal [4]. Recent, preliminary clinical studies have suggested that virucidal activity of oral rinses may occur in vivo against SARS-CoV2 [6, 7].

Since only one of the *in vitro* studies has used the SARS-CoV-2 pathogen to date, here, we extended this work by testing the virucidal activity against SARS-CoV-2 of a range of mouthwashes including CPC (0.05-0.1%w/v Dentyl Dual Action, 0.05-0.1% w/v Dentyl Fresh Protect, 0.10% w/v SCD Max) ethanol/essential oils (Listerine Cool Mint, 21.7% v/v ethanol), ethanol/ethyl lauroyl arginate (Listerine Advanced Gum Treatment, 23% v/v ethanol), chlorhexidine (0.2% w/v; Corsodyl) and povidone iodine (0.5% w/v; Videne). We found that three products had sufficient activity to pass EN14476 against SARS-CoV2. We also investigated the contribution of ethanol to the observed virucidal activity, to inform future studies as to which products are most likely to provide the greatest benefit against SARS-CoV2, and to provide information as to how future virucidal formulations might be optimised.

## Methods

Virucidal assays utilised VeroE6, a gift from the University of Glasgow/MRC Centre for Virology, UK. The England2 strain of SARS-CoV2 was provided by Public Health England, and amplified in VeroE6 cells before being harvested from the supernatant. All cells were grown in DMEM containing 10 % (v/v) FCS, and incubated at 37 °C in 5 % CO_2_. Virucidal activity of mouthwash was studied in media containing 100 μL mucin type I-S, 25 μL BSA Fraction V, and 35 μL yeast extract to mimic oral secretions. 100 μL of this mixture was added to 100 μL of virus suspension, and 800 μL of the test-product added. After 30 seconds, virucidal activity was neutralised by 10-fold serial dilution in ice-cold DMEM (containing 10% FCS). Alternatively, in a modification of the methods of Mesiter [3], virus was purified by size-exclusion chromatography (SEC) to prevent direct cytotoxic effects of the products on the cell monolayer; 100 μL of the mixture was added to a microspin S-400 HR column, and centrifuged for 2min at 700 × *g*. A 10-fold serial dilution was then made of the flow-through in DMEM containing 10% FCS. In a further modification to the methods of Meister *et al* [3], we titrated virus onto VeroE6 cells transduced with Lentivirus vectors expressing ACE2 and TMPRSS2 and drug selected, to enhance virus entry (>1-log), generating a more sensitive test for virucidal activity. Titrations were performed by plaque assay; serial dilutions were used to infect VeroE6/ACE2/TMPRSS2 cells for 1 h. Following this, cells were overlaid with DMEM containing 2 % FCS, and 1.2 % Avicel^®^. After 72 h, the overlay was removed, and the monolayer washed and fixed with 100% methanol. Monolayers were stained with a solution of 2 5% (v/v) methanol and 0.5 % (w/v) Crystal Violet, then washed with water, and plaques were enumerated.

## Results

In initial experiments, we examined the effect of mouthwashes on the VeroE6/ACE2/TMPRSS2 monolayers used to detect live virus, following serial dilution in DMEM. Four of the seven products demonstrated toxicity to the monolayer, which was not eliminated until they were diluted at least 100-1000-fold, limiting the sensitivity of the assay to measure residual infectivity. We therefore used size-exclusion chromatography (SEC) to rapidly purify virus away from the products. When virus was purified on S-400 HR microspin columns, only minimal loss of infectivity was observed (Fig 1A), however toxicity from the mouthwashes against the cell monolayer was virtually eliminated (Fig 1B). SEC was therefore used for all assays from this point forwards. This approach also ensures that the activity of the mouthwashes against the virus was rapidly stopped after the desired co-incubation time. In comparison, our results suggest that ‘stopping’ the reaction by serial 10-fold dilution would leave sufficient mouthwash activity to have continued biological activity against the virus. Combined with the use of VeroE6/ACE2/TMPRSS2 cells for titration, which SARS-COV2 enters >1log more efficiently than parental VeroE6, the assay was capable of detecting a 5-log10 decrease in virus titre. This is more than sufficient to detect the 4-log10 reduction in activity specified by EN14476.

**Figure 1.**
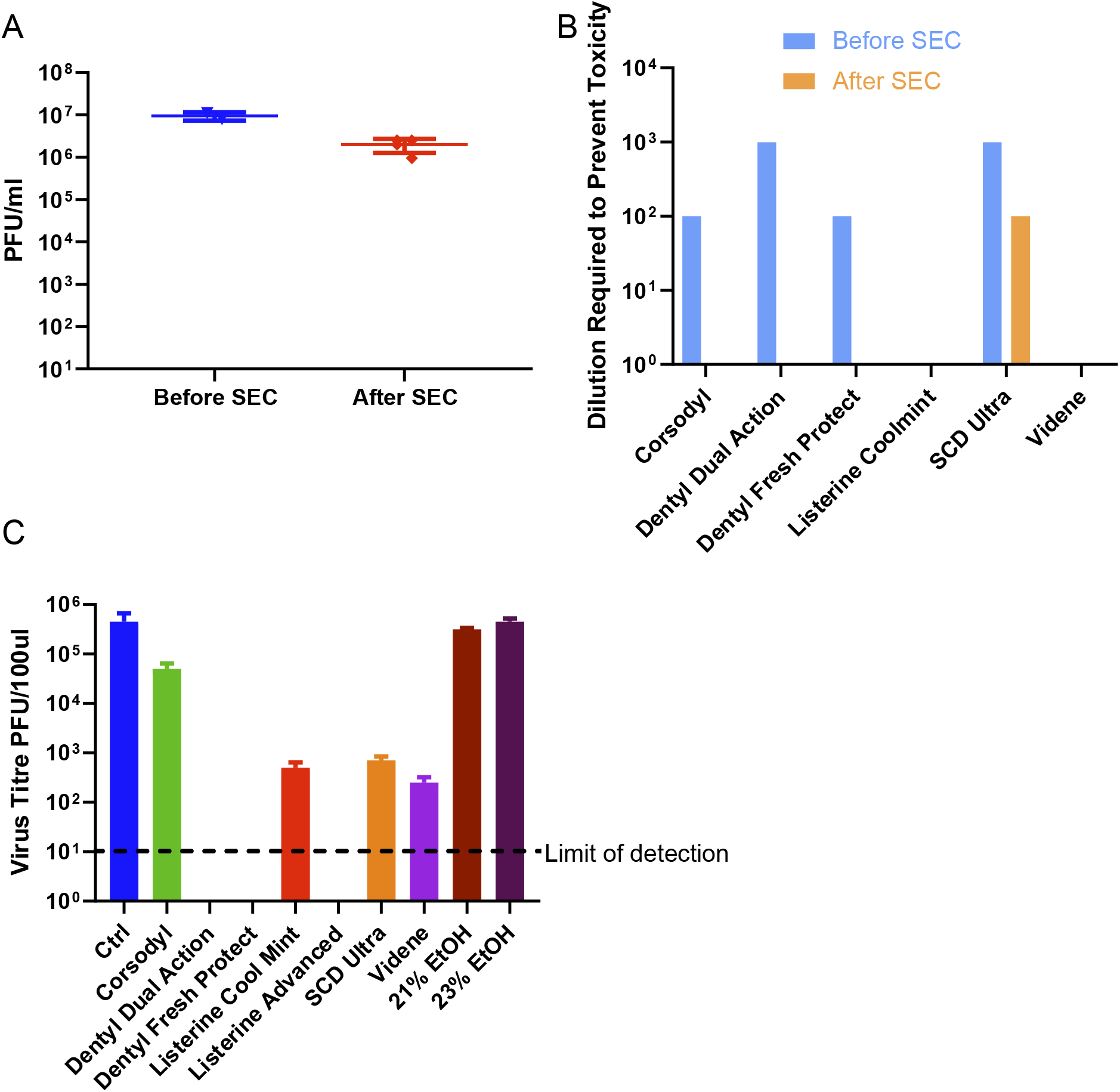
(A) 100μl virus was purified through a S-400 HR spin column, and live virus measured by plaque assay on VeroE6/ACE2/TMPRSS2. Only minimal loss of virus titre was observed. (B) Mouthwashes were mixed with DMEM (instead of virus) and synthetic salivary secretions, then 100μl of the mixture was purified through a S-400 HR spin column, diluted by serial 10-fold dilution in DMEM/10, and titrated onto VeroE6/ACE2/TMPRSS2. After 72h, overlays were removed and monolayers were fixed and stained with crystal violet, then toxicity was scored based on visual inspection of monolayer integrity. (C) Virus was mixed with synthetic salivary secretions and mouthwash, then purified by SEC after 30 seconds, before being titrated by plaque assay on VeroE6/ACE2/TMPRSS2.

Having optimised a sensitive protocol for the detection of virucidal activity, we tested the ability of a wide range of commercially available mouthwash formulations (Table 1) to reduce virus infectivity, after a 30-second treatment. The mouthwashes demonstrated a wide spectrum of inactivation ability (Fig 1C). Two Dentyl mouthwashes containing CPC, and Listerine Advanced (23 % ethanol) with ethyl lauroyl arginate (LAE), a cationic surfactant, eradicated the virus completely, giving >5-log10 reduction in viral titres. A moderate effect (~3-log fold reduction) was seen with the iodine containing product (Videne), SCD Max (CPC and Sodium Citric Acid), and mouthwash containing 21 % v/v alcohol with essential oils (Listerine Cool Mint) (Fig 1C). Ethanol alone at <23 % had no effect on virus infectivity, thus the inclusion of essential oils (Listerine Cool Mint) or LAE (Listerine Advanced) appears to be required for optimal efficacy. Lastly, chlorhexidine was relatively inactive (<2 log fold reduction).

**Table 1.**

| Product name | Active Ingredients |
| --- | --- |
| **Corsodyl** | 7 % (v/v) ethanol, 0.2 % (w/v) chlorhexidine<br>Other active ingredients: peppermint oil |
| **Dentyl Dual Action** | 0.05 %-0.1 % (v/v) cetylpyridinium chloride (CPC),<br>Other active ingredients: isopropyl myrisate, Mentha Arvensis extract |
| **Dentyl Fresh Protect** | 0.05 %-0.1 % (v/v) cetylpyridinium ehloride (CPC)<br>Other active ingredients: xylitol |
| **Listerine Cool Mint** | 21 % (v/v) ethanol<br>Other active ingredients: thymol 0.064 %, eucalyptol 0.092 %, methyl salicylate 0.060 % and menthol 0.042 % |
| **Listerine Advanced Gum Treatment** | 23 % (v/v) ethanol,<br>Other active ingredients: ethyl lauroyl arginate HCl (LAE) 0.147% w/w. |
| **SCD Max** | 0.07-0.1 % (v/v) cetylpyridinium chloride (CPC) and sodium citric acid 0.05 %<br>Other active ingredients: sodium monofluorophosphate. |
| **Videne** | 7.5% iodinated povidone equivalent to 8.25 mg/ml iodine |

## Discussion

Our data, using an assay that enabled us to detect up to 5-log10 reduction in SARS-CoV2 activity, further support the accumulating evidence for high virucidal activity by widely-available mouthwashes *in vitro*, against SARS-CoV2 and the related HCoV-229E coronavirus strain [3-5]. Here, three products which contained either (i) 0.07-0.1 % CPC (Dentyl Dual Action, Dentyl Fresh Protect) or (ii) 23 % ethanol with LAE (Listerine Advanced) provided the greatest level of inactivation, surpassing the level required for EN14476.

The ability of CPC-containing mouthwashes to reduce viral infectivity is in line with studies using other enveloped viruses; similar microemulsion formulations of CPC (with a similar isopropyl myristate microemulsion as Dentyl Dual Action) have been demonstrated to exhibit both inherent antimicrobial activity [8] and specific activity against Herpes Simplex virus [9], and other mouthwash formulations containing 0.07 % CPC have been reported to reduce infectivity of seasonal coronaviruses by 3-4-log10 [4]. In our study, SARS-CoV2 was even more sensitive to the CPC-containing mouthwashes than in these previous reports using hCoV-229E; this may reflect the cumulative activity of additional components in these formulations. The SCD Max (containing CPC at a higher concentration) only resulted in a 3-log10 reduction in infectivity, and suggests that the exact formulation is important; thus, individual mouthwash formulations should be empirically tested for antiviral activity, rather than basing decisions on the ‘major’ antimicrobial component.

Listerine (Cool Mint, Ultra, Antiseptic formulations) was recently shown by others to have virucidal activity towards both SARS-CoV2 or HCoV-229E. Meyers *et al* showed >4 log10 reduction in titres for Listerine Antiseptic and 3-4 log10 reductions for Listerine Ultra against HCoV-229E [4], while Meister *et al* found either >2 or >3 log10 reductions for Listerine Cool Mint against 3 separate SARS-CoV2 strains isolated from patients [3, 4]. Here, the greater sensitivity of our assay revealed that while Listerine Cool Mint reduced infectivity by 3-logs, Listerine Advanced was superior, and was capable of totally inactivating SARS-CoV2, reducing infectivity by >5-log10. As with the CPC-containing products, the importance of formulation was evident in alcohol-containing preparations; alcohol alone at the same concentration as in Listerine Cool Mint or Advanced had minimal impact, indicating that the essential oils and LAE in these formulations significantly augment antiviral activity. Whilst poorly-defined chemically, the antiviral activity of essential oils present in the Dentyl Products and Listerine products (a mixture of plant-derived monoterpenes and phytoretinoids) has been previously extensively described [10], albeit not in the context of SARS-CoV2. Moreover, the addition of the cationic surfactant LAE, which is recognised to exhibit antiviral activity *in vitro* [11], resulted in a >2-log increase in activity compared to the Listerine Cool Mint product; resulting in a product that gave an equal level of kill to the CPC-containing products.

Two very preliminary studies using very small numbers of patients with COVID19 have suggested that mouthwashes including PVP-I [6] and chlorhexidine [7] may reduce SARS-CoV2 loads *in vivo*. However, these studies used qPCR, which does not determine whether the detected virus is infectious, the numbers of patients studied was extremely low, and the impact was variable. While these studies demonstrate a potential for reducing the level of SARS-CoV2 in the oropharynx, it is important to note that *in vivo* studies are currently lacking into how effective such an approach might be *in vivo* at reducing viral titre and transmission. It is critical to determine how quickly virus shedding from actively infected cells in both the upper and lower respiratory tract replenishes live virus in the oral cavity after treatment. Our *in vitro* data identifies products with high activity, and indicates that investigating the duration of their effects *in vivo* against live virus load, and defining potential effects on reducing the risk of virus exposure within the clinical setting (for example when performing clinical examinations of the oropharynx, or visiting vulnerable elderly populations/patients), in well-designed, randomised controlled trials is warranted. Importantly the anti-viral mechanisms of action for oral rinses are dependent on the lipid composition of the viral envelope and its sensitivity to surfactants and membrane disrupting agents. Since this membrane derives from host cell membranes, it is unlikely to be altered by virus mutation.

